# Sequence-Based Therapeutic Peptide Classification with Augmented Negative Sampling

**DOI:** 10.64898/2026.06.07.730473

**Authors:** Roman Ellerbrock, Alessio Valentini, Alexander C. Paul, Swagatam Mukhopadhyay, Michael R. Perelshtein

## Abstract

Therapeutic peptides offer high target specificity, low toxicity, and the ability to modulate protein-protein interactions, yet experimental functional characterization remains costly and slow. Computational prediction of therapeutic function directly from sequence could accelerate peptide screening and enable generative design pipelines, but requires reliable discrimination between therapeutic and non-therapeutic peptides. Existing multi-label predictors cover few functions, rely on limited datasets, and exhibit high False Positive Rates (FPRs), limiting their practical utility.

We present a lightweight CNN classifier trained on the most comprehensive therapeutic peptide database to date (54,655 peptides, 48 functional categories). A key contribution is a statistically motivated negative sampling strategy using Markov models to generate diverse *synthetic decoys* at multiple difficulty levels. When evaluated on this controlled decoy benchmark, the FPR is reduced from over 60% for previous models to 2.1% for our approach. On positive therapeutic samples, our fine-tuned five-model ensemble achieves 79.9% Micro F1 and 54.6% Macro F1 while requiring only amino acid sequences as inputs.

Analysis using a sparse L1-constrained variant of our model shows that convolutional filters capture conserved functional motifs and statistically improbable non-therapeutic patterns, with downstream layers combining these signals, providing mechanistic evidence that the network learns biologically meaningful structure. On an external generalization benchmark derived from TPpred-LE, our model achieves 55.3% Micro F1 and 38.6% Macro F1 on the 12 shared labels, close to the benchmark-specific baseline (57.9%/38.1%), while retaining substantially broader therapeutic label coverage. Code and models will be made available at https://github.com/terra-quantum-public/tq-therapep-ai.

## 1 Introduction

Therapeutic peptides have emerged as a promising class of drug candidates, with the global market projected to exceed $50 billion by 2030 [Muttenthaler et al., 2021]. Their appeal stems from a unique combination of properties: higher target specificity than small molecules, lower immunogenicity than antibodies, and the ability to modulate protein-protein interactions – targets historically considered “undruggable” by conventional pharmaceuticals [Fosgerau and Hoffmann, 2015]. Applications span antimicrobial, anticancer, antiviral, antihypertensive, and immunomodulatory therapies [Xiao et al., 2025], with antimicrobial peptides receiving particular attention amid the growing antibiotic resistance crisis [Mahlapuu et al., 2016].

Despite this potential, identifying therapeutic peptides remains a bottleneck. Experimental characterization of peptide function is costly and time-consuming: a single peptide candidate may require weeks of synthesis, purification, and bioactivity assays, while the average cost of bringing a drug to market exceeds $2 billion [DiMasi et al., 2016]. Even with curated databases of tens of thousands of known therapeutic peptides, experimental validation of novel candidates or repurposing of existing sequences for new functions remains prohibitively slow. Computational prediction of therapeutic function directly from amino acid sequences could dramatically accelerate drug discovery by prioritizing candidates for experimental validation, potentially reducing development timelines and costs significantly.

A key advantage of sequence-based prediction is that it requires no structural information. While protein structure can inform function, structure determination remains expensive and not always feasible for short peptides that may lack stable folds. In contrast, amino acid sequences are readily available from genomic data, peptide databases, and rational design pipelines. An accurate sequence-based classifier would enable high-throughput virtual screening of candidate peptides without the bottleneck of structure prediction or determination. More importantly, it is not clear that protein Language Models are accurate in predicting the low-energy conformations of therapeutic peptides, which are often chemically-modified to improve pharmacological properties like bio-availability and binding affinity. The introduction of non-canonical residues, macro-cyclization etc. makes therapeutic peptides chemically distinct from proteins [Quartararo et al., 2020, Fetse et al., 2023], and very recent work reiterates this sentiment [Feller et al., 2026]. Predicting therapeutic peptide function, however, presents several fundamental machine learning challenges. First, it is inherently a *multi-label* problem: a single peptide may exhibit multiple therapeutic activities (e.g., both antimicrobial and anticancer). Second, the label distribution is severely *imbalanced* : in the recently published therapeutic peptide database (TheraPepDB)[Xiao et al., 2025], antibacterial peptides number in the tens of thousands while rare functions like Quorum Sensing have fewer than 100 examples. Third, and perhaps most critically, classifiers must avoid *false positives* – incorrectly predicting therapeutic activity for inactive sequences.

Most existing models focus almost exclusively on distinguishing between therapeutic classes, while the negative class is often poorly defined, weakly represented, or entirely absent. As a consequence, reported performance metrics primarily reflect the ability to separate therapeutic subclasses rather than the ability to reject non-therapeutic sequences. In real screening scenarios, however, the overwhelming majority of candidate peptides are non-therapeutic. Even a modest FPR can therefore lead to an impractically large number of experimentally wasted candidates, severely limiting the usefulness of computational predictors.

Existing computational approaches to therapeutic peptide prediction fall into two categories. The first comprises specialized binary classifiers for individual functions: tools like iAMP-2L for antimicrobial peptides [Xiao et al., 2013], AntiCP for anticancer peptides [Tyagi et al., 2013], and AVPpred for antiviral peptides [Thakur et al., 2012]. While effective for their specific targets, these tools cannot capture the multi-functional nature of therapeutic peptides and require users to run multiple separate predictions. The second category comprises multi-label classifiers that predict multiple functions simultaneously. TPpred-LE [Lv et al., 2023a][Lv et al., 2023b] employs a transformer encoder-decoder architecture with label embeddings to capture correlations between therapeutic functions, achieving 57.9% Micro F1 on a 15-label benchmark with approximately 23 million parameters. TPpred-CMvL [Yan et al., 2024] extends this approach using contrastive multi-view learning with TAPE protein embeddings and position-specific scoring matrices (PSSM), achieving 54.0% example accuracy. Most recently, PLMCCL-TP [Lin and Tuo, 2025] combines pre-trained protein language models (ProtBERT) with Gaussian mixture model clustering and contrastive learning, achieving 72.8% precision on a 21-label benchmark. A very recent paper introduced a rather large SMILES-based chemical language model, with augmenting peptide chemical space learning with small molecule and lipid datasets [Feller et al., 2026].

Despite these advances, current multi-label methods face several limitations. First, they cover only 12–21 therapeutic functions, leaving dozens of clinically relevant categories unpredicted. Second, they rely on large transformer-based architectures with tens of millions of parameters, which may be unnecessary for short peptide sequences where local motifs dominate over long-range dependencies. Third, many require external features such as PSSM profiles or pre-trained protein language model embeddings, increasing computational complexity. Finally, and most critically for real-world screening, existing methods exhibit high FPRs when evaluated against realistic negative sequences, limiting their practical utility in large peptide libraries dominated by inactive candidates.

We hypothesize that therapeutic peptide function is primarily determined by short sequence motifs – a hypothesis supported by known functional signatures such as the YGGF enkephalin motif [Hughes et al., 1975] in opioid peptides and the KLAK pro-apoptotic helix [Ellerby et al., 1999] in anticancer peptides. Convolutional Neural Networks (CNNs) are well-suited to detect such local patterns through their translation-invariant filters.

In this work, we present a sequence-based CNN classifier for multi-label therapeutic peptide function prediction. Our main contributions are:

1. **Augmented negative sampling using synthetic decoy sequences**. One of our major contributions is leveraging *synthetic decoy* sequences to create a negative class, especially for classes of peptides for which data is biased and small in size. We show that our approach reduces false positive rate in our classifier. These *synthetic decoy* sequences are *statistically similar* to therapeutic peptides but are highly unlikely to have functional motifs. We posit that this forces the classifier to learn discriminative features rather than superficial compositional biases. We create these synthetic decoy sequences using Markov models at multiple difficulty levels (uniform random, global Markov, position-dependent Markov, and class-frequency sampling). This approach substantially reduces FPRs compared to naive random sampling.
2. **Efficient architecture**. We provide two model variants: a lightweight CNN (1M parameters) suitable for rapid screening, and a fine-tuned 5-model ensemble (15M parameters) that achieves 79.9% Micro F1 and 54.6% Macro F1 on positive therapeutic samples, and 78.9% Micro F1 and 54.4% Macro F1 on the full held-out test set. Both models only require amino acid sequences as input.
3. **Competitive external performance**. On the TPpred-LE benchmark, after removing all overlapping sequences from our training data, our model achieves 55.3% Micro F1 and 38.6% Macro F1, comparable to TPpred-LE trained on its native dataset (57.9% and 38.1%) while retaining substantially broader therapeutic label coverage.
4. **Biological interpretability and motif analysis**. We analyze the learned representations using multiple sequence alignments and sparse L1-constrained model variants, showing that convolutional filters capture conserved functional motifs as well as statistically improbable non-therapeutic patterns. The downstream layers combine these signals to reproduce biologically meaningful logic, providing mechanistic evidence that the model learns functionally relevant sequence features rather than dataset-specific artifacts.

The remainder of this paper is organized as follows. Section 2 describes our dataset, negative sampling strategy, model architecture, and training procedure. Section 3 describes the experimental setup and evaluation metrics. Section 4 presents our results and discussion, including comparisons with prior methods, external benchmark evaluation, architecture analysis, error patterns, and model interpretability. Section 5 concludes with future directions.

## 2 Methods

### 2.1 Dataset

#### 2.1.1 Therapeutic Peptides

We use the comprehensive therapeutic peptide dataset of Xiao et al. [2025], comprising 58,583 therapeutic peptides. From this resource, we selected 54,655 natural therapeutic peptides annotated with 48 therapeutic function labels plus one global label indicating whether the sequence has therapeutic function. Hyperparameter tuning was performed exclusively on this natural-peptide subset.

During model fine-tuning, we additionally incorporated 1,000 non-standard peptides containing ambiguous amino acids arising from mass-spectrometry indistinguishability (e.g., isoleucine and leucine, glutamine and lysine) in order to improve sampling of rare functional classes. Empirically, their inclusion led to a noticeable increase in macro F1. We therefore retain these sequences, motivated by the assumption that the minor noise introduced by random resolution of a small number of ambiguous residues is outweighed by the benefit of improved class coverage compared to excluding these peptides entirely. Peptide lengths range from 2 to 50 Amino Acids (AA), and a single outlier with 74 AA, with a mean of 18.2 AA (Figure 1). The structural annotations indicate that many therapeutic peptides in this dataset adopt regular secondary structures, particularly helical and *β*-sheet-like conformations. Because such secondary-structure propensities are often predictable from sequence-derived features, this supports the view that local sequence motifs capture substantial functional signal.

**Figure 1:**
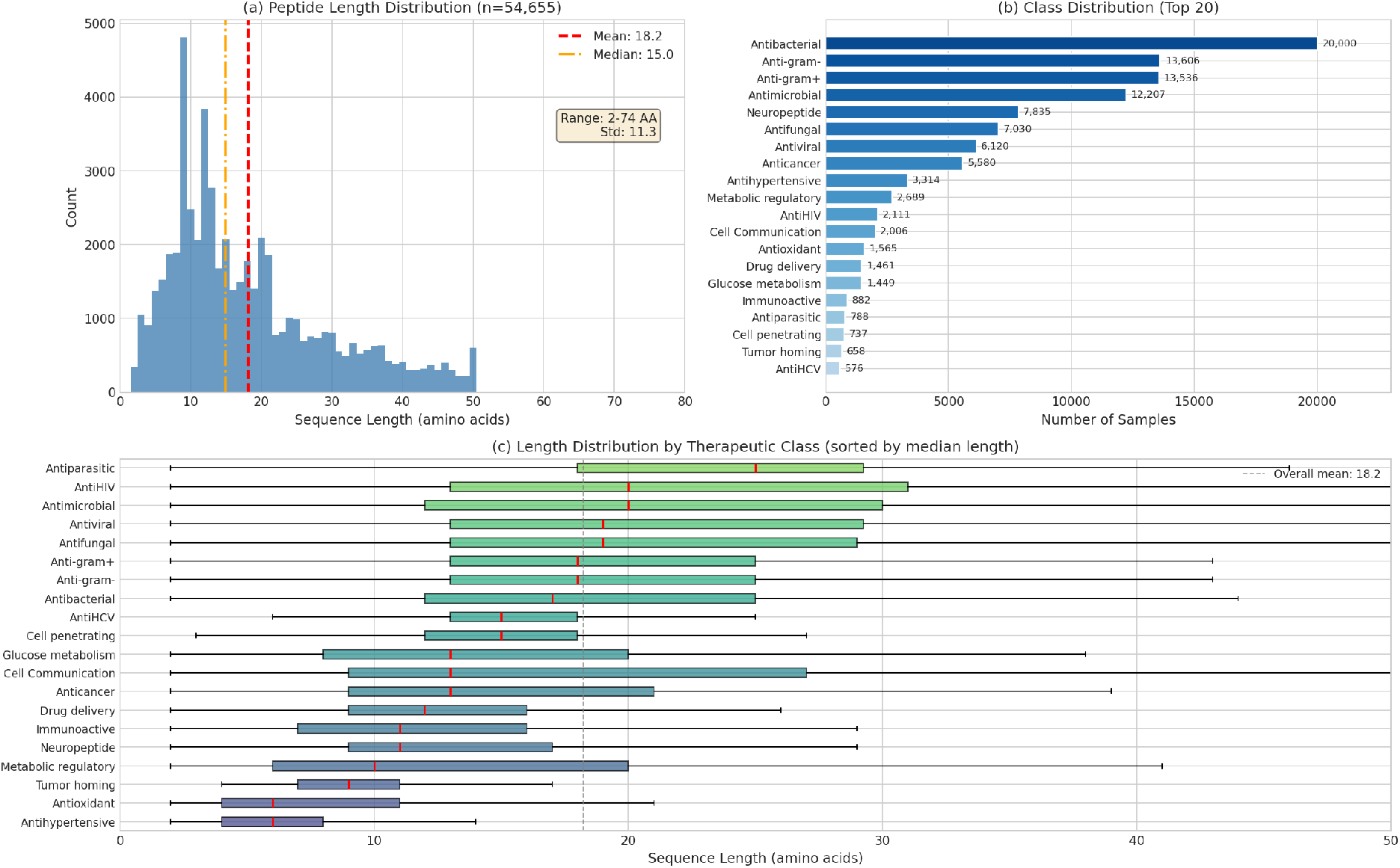
Dataset Overview. (a) Peptide length distribution showing predominantly short sequences (mean 18.2 AA, median 15 AA). (b) Class imbalance across therapeutic functions –Antibacterial peptides dominate (20,000 samples) while rare functions have fewer than 100 samples. (c) Length distribution by therapeutic class reveals functional signatures: Antihypertensive peptides are notably short (median ∼6 AA), while Antimicrobial peptides are longer (median ∼ 20 AA). Negative decoys are generated to prevent our model from learning simple correlation patterns.

The dataset exhibits significant class imbalance: Antibacterial peptides constitute the largest class (20,000 samples), while rare functions like quorum sensing have fewer than 100 samples. Different therapeutic classes show distinct amino acid preferences (Figure 2): antimicrobial peptides are enriched in cationic residues (K, R), while antihypertensive peptides favor proline (P). These class-specific compositional signatures extend to dipeptide motifs (Figure 3), providing discriminative features for classification.

**Figure 2:**
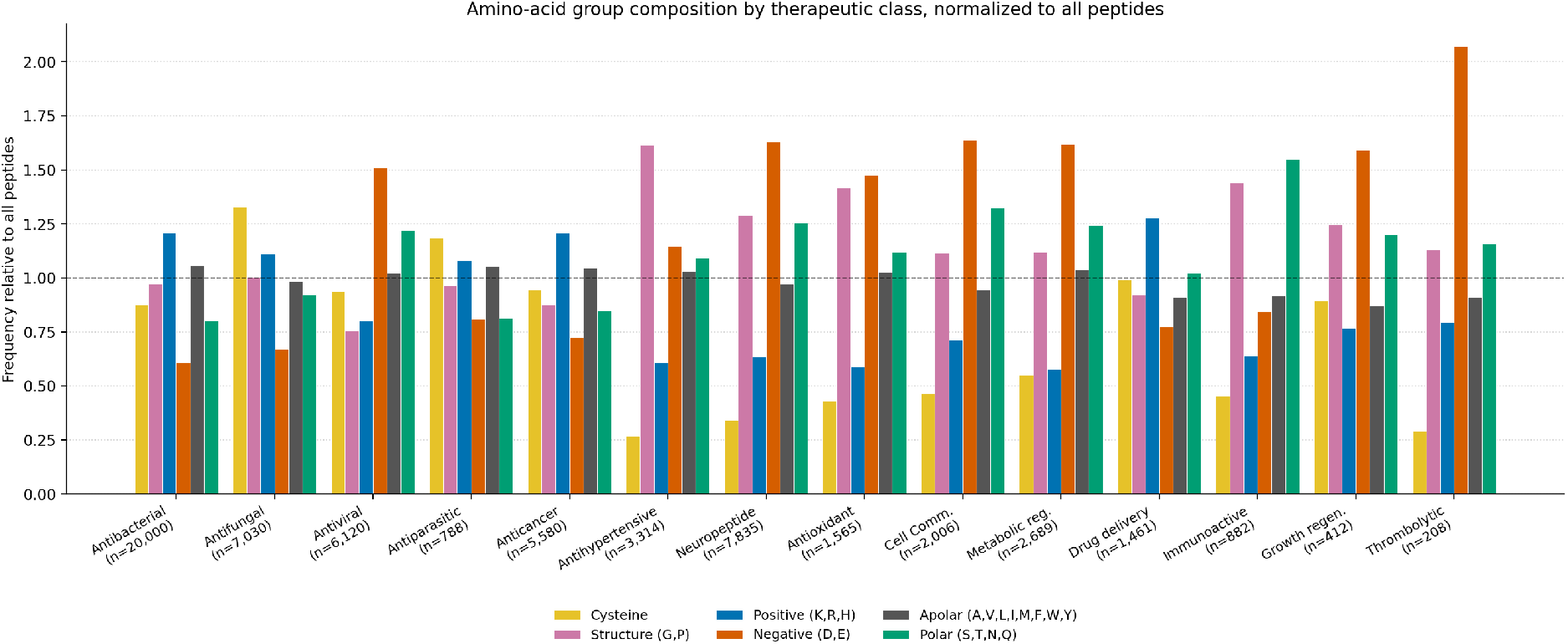
Amino-Acid Group Composition by Therapeutic Class. Residue composition of six biochemical groups—cysteine, structure-modifying (G,P), positive (K,R,H), negative (D,E), apolar (A,V,L,I,M,F,W,Y), and polar (S,T,N,Q)—normalized to their dataset-wide frequency (dashed line). Each class exhibits a distinctive biochemical signature consistent with its mechanism of action: antimicrobial classes (antibacterial, antifungal, antiviral, antiparasitic) and drug-delivery peptides are enriched in positive charge (∼ 1.2–1.5×) and depleted in negative charge, consistent with electrostatic attraction to anionic membranes. Antifungal and antiparasitic peptides additionally show elevated cysteine (∼1.2–1.35×), reflecting disulfide-stabilized scaffolds such as defensins. Antihypertensive and neuropeptides are enriched in structure-modifying residues (G, P; up to ∼ 1.6 ×), in line with the proline-rich ACE-inhibitor signature and the flexible turn motifs characteristic of neuropeptides. While these differing compositional biases are informative for class discrimination, biological function requires more than simple compositional imbalances - sequence order and local motif structure also matter

**Figure 3:**
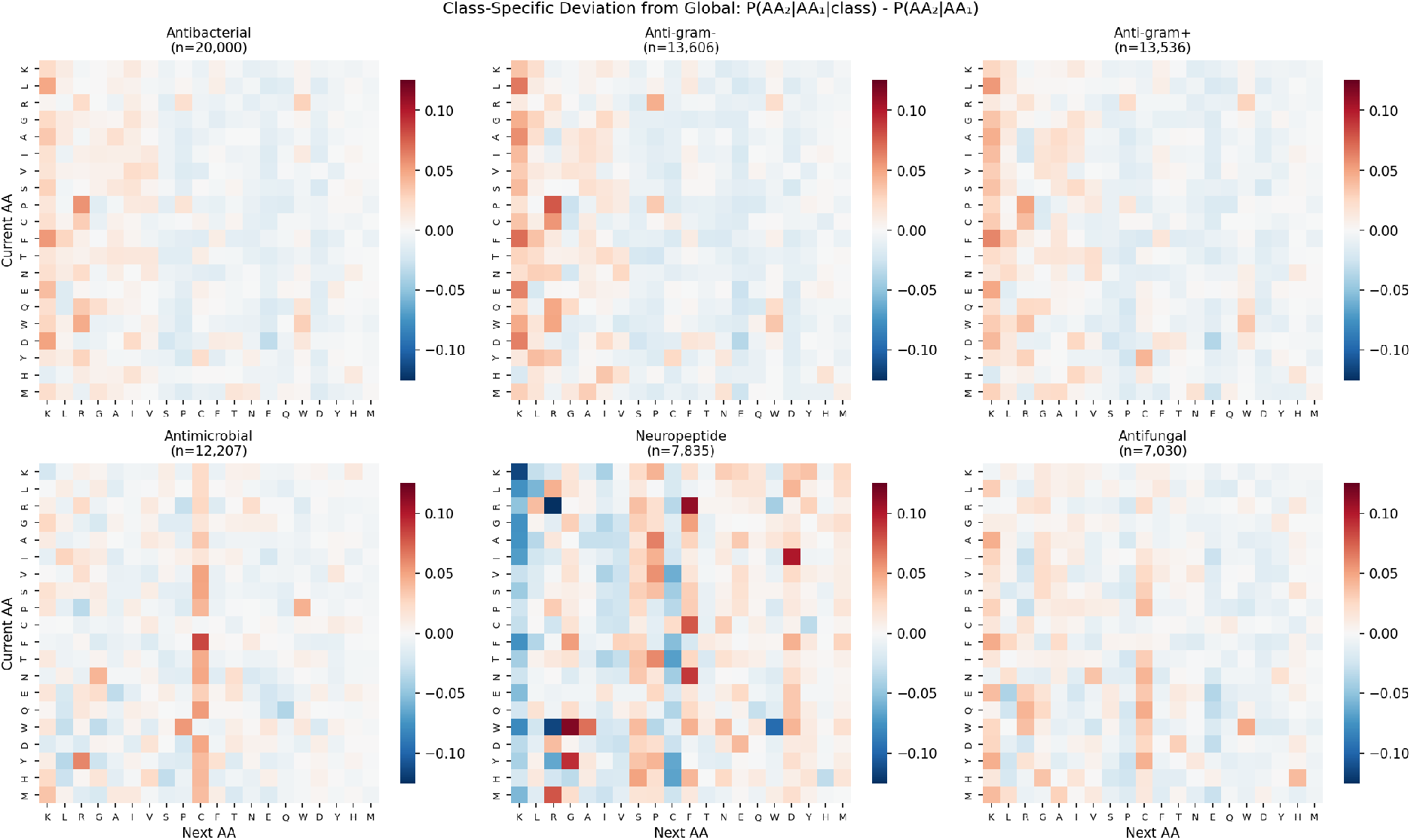
Class-Specific Dipeptide Signatures. Difference in dipeptide frequencies between each therapeutic class and the global average. Positive values (red) indicate enrichment; negative values (blue) indicate depletion. These distinctive local sequence motifs are captured by our CNN’s convolutional filters. Please note that the global average is dominated by antimicrobial peptides which make a large portion of the dataset.

Multiple sequence alignment (MSA) analysis reveals biologically validated motifs that correlate with therapeutic function. Opioid and analgesic peptides share the conserved N-terminal motif YGGF (enkephalin signature, found in 41% and 14% respectively), which binds to opioid receptors [Hughes et al., 1975]. Anticancer peptides across multiple cancer types (breast, colon, lung, skin) are enriched in the KLAK motif (14–30%), a pro-apoptotic amphipathic helix that disrupts mitochondrial membranes [Ellerby et al., 1999]. Cell-penetrating peptides show the cationic RKKR motif (6%) [Frankel and Pabo, 1988, Green and Loewenstein, 1988], while tumor-homing peptides contain the integrin-binding RGD sequence (5–9%) [Pierschbacher and Ruoslahti, 1984]. Antiviral peptides exhibit virus-specific signatures: AntiHIV peptides are enriched in QQEK (12.7%), AntiHCV in poly-lysine KKKK (26.6%), and AntiHSV in disulfide-forming RCIC (15%) [Urmi et al., 2023]. These functionally validated motifs, spanning 4–8 AA, align well with our CNN’s convolutional kernel sizes (3, 5, 7, 9), supporting the architectural choice for capturing therapeutically relevant sequence patterns while long kernels (7, 9) can pick up on statistical properties.

#### 2.1.2 Synthetic Decoy Generation

The dataset analysis in Section 2.1.1 reveals strong class-specific compositional signatures: distinct length distributions (Figure 1c), amino acid preferences (Figure 2), and dipeptide frequencies (Figure 3). These differences are pronounced enough that a classifier could plausibly achieve high accuracy by learning compositional shortcuts alone, without capturing the sequence-level features that actually determine therapeutic function. Such a model would then misclassify any peptide-like sequence sharing similar statistics as therapeutic — catastrophic for screening, where most candidates are non-therapeutic.

We address this by introducing synthetic decoys that match the compositional statistics of therapeutic peptides while lacking their functional sequence structure. By forcing the model to discriminate against decoys with matching amino acid frequencies, dipeptide transitions, and positional biases, we eliminate compositional shortcuts and require the model to learn higher-order patterns like local motifs and ordered residue arrangements that genuinely distinguish therapeutic function from peptide-like composition. The four-tier strategy below provides a gradient of difficulty that progressively removes each level of compositional similarity.

This strategy inevitably introduces false negatives, that is, decoys that may in fact exhibit therapeutic activity. We deliberately accept this trade-off: truly therapeutic peptides constitute a vanishingly small fraction of sequence space, and in screening workflows, false positives translate directly into wasted synthesis effort. The necessity of this approach is confirmed empirically: previously published predictors classify our decoys as therapeutic in 60–70% of cases, frequently within rare classes such as anti-HIV or anticancer peptides, confirming that existing models have learned compositional shortcuts rather than functional sequence features.

Our approach employs four complementary generation strategies, each targeting different aspects of sequence composition:

1. **Uniform random**: Each amino acid is sampled independently with equal probability *P* (*AA*) = 1*/*20. These sequences lack any compositional bias and serve as “easy” negatives, ensuring the model learns basic amino acid preferences of therapeutic peptides.
2. **Global Markov**: First-order Markov chain with transition probabilities *P* (*AA*_*i*+1_|*AA*_*i*_) estimated from all therapeutic sequences. These decoys preserve dipeptide frequencies but lack position-specific or class-specific patterns, representing “medium” difficulty negatives.
3. **Position-dependent Markov**: Transition probabilities conditioned on sequence position, *P* (*AA*_*i*+1_ | *AA*_*i*_, *i*), capturing the observation that certain positions (e.g., N-terminus, C-terminus) have distinct compositional biases. These “hard” negatives mimic positional preferences while lacking functional motifs.
4. **Class-frequency sampling**: Amino acid frequencies estimated separately for each therapeutic class, *P* (*AA* |*class*). Sequences are generated to match the composition of specific functional categories (e.g., antimicrobial, anticancer) without preserving the sequential patterns that confer activity. These represent the “hardest” negatives, forcing the model to learn beyond simple composition.

We generated decoy sequences to obtain an overall 1:2 positive-to-negative ratio, with equal contributions from four negative-sampling strategies. This class balance reflects the intended learning setting, in which non-therapeutic sequences are more frequent than therapeutic ones and, in the absence of informative features, the statistically favored prediction is therefore non-therapeutic. To prevent the model from exploiting sequence length as a trivial discriminative feature, decoy lengths were sampled from the empirical length distribution of therapeutic peptides. This diverse negative-sampling scheme exposes the model to a spectrum of decoy difficulty, ranging from compositionally simple random sequences to more sophisticated mimics that differ from therapeutic peptides only in subtle sequential patterns.

We also evaluated mutation- and shuffle-based decoy generation. In the mutation-based approach, approximately 50% of the AAs in therapeutic sequences were randomly substituted, whereas the shuffle-based strategy preserved amino-acid composition while randomizing sequence order. However, both approaches substantially reduced model performance on positive samples. This likely occurred because, for some therapeutic categories, activity may be retained despite extensive mutation or shuffling, particularly when key functional motifs remain partially intact. As a result, these strategies may generate false negatives that are still recognized as active by the model, thereby weakening the distinction between therapeutic and non-therapeutic classes.

We emphasize that this decoy generation strategy represents a first step toward reducing false positives, not a complete solution. The fundamental challenge remains: synthetic decoys, however sophisticated, cannot fully capture the complexity of the biological sequence space. Future work could incorporate experimentally validated inactive sequences, leverage structural information to generate physically implausible decoys, or employ adversarial generation methods that specifically target model weaknesses. A particularly promising direction is active learning: using the trained classifier to identify sequences near the decision boundary, then iteratively generating targeted decoys that refine the distinction between therapeutically active and inactive peptides. Nevertheless, our multi-strategy approach provides a principled foundation that demonstrably improves model robustness compared to naive random sampling, as validated in our ablation studies (Section 4).

### 2.2 Model Architecture

Based on a systematic evaluation of alternative sequence modeling approaches (see subsequent subsections), we selected a lightweight two-layer CNN architecture for therapeutic peptide clas-sification (Figure 4). This choice was guided by comparative benchmarking and by our working hypothesis that therapeutic function in short peptides is predominantly governed by local sequence motifs rather than long-range dependencies. This hypothesis is consistent with the conserved motif patterns identified in our MSA analysis. In practice, the two-layer CNN provided the best trade-off between predictive performance, parameter efficiency, and robustness across functional classes, while avoiding unnecessary architectural complexity for sequences of length 10–50 residues.

**Figure 4:**
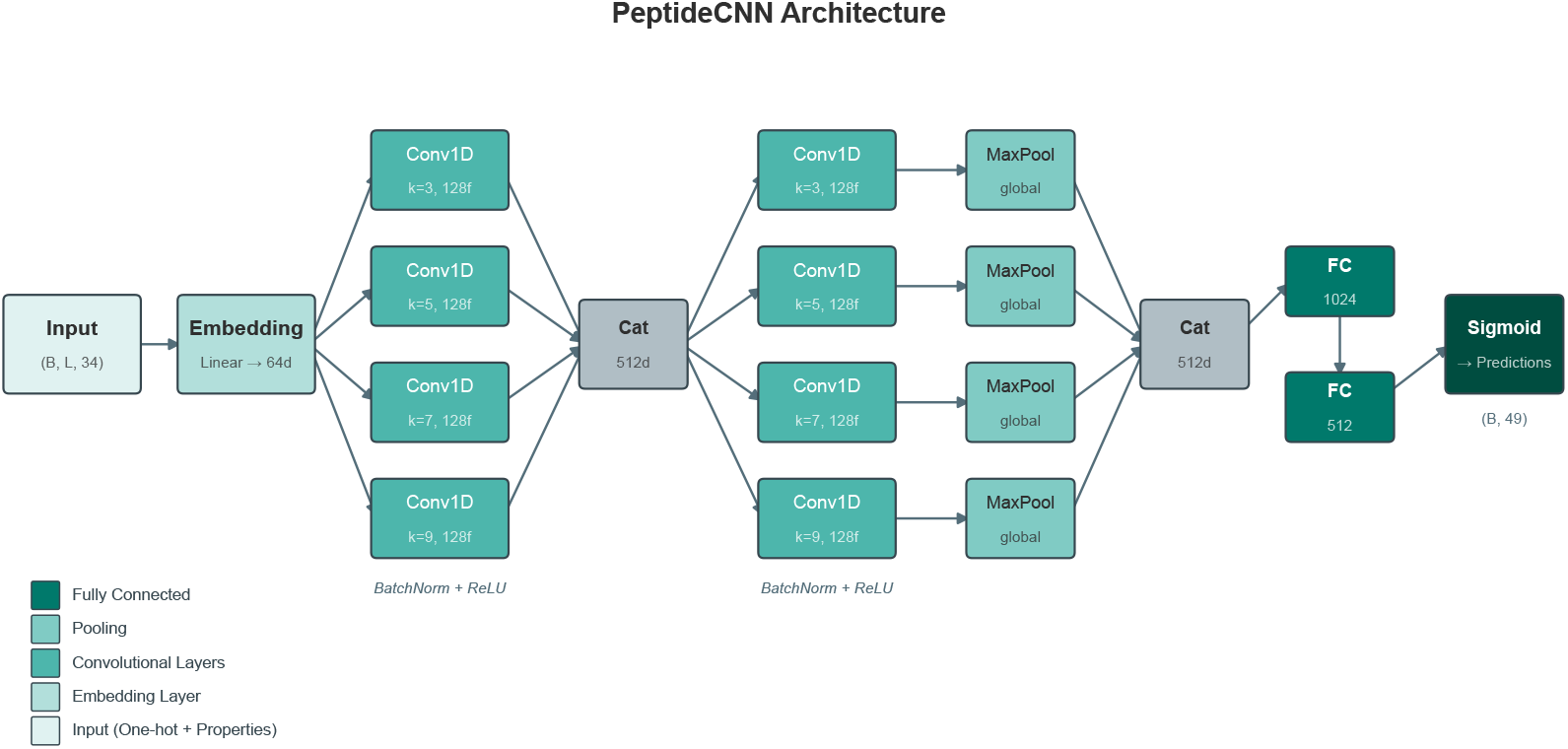
Model Architecture. Our two-layer CNN uses parallel convolutions with kernel sizes (3, 5, 7, 9) to capture local sequence motifs at multiple scales. The first convolutional block extracts basic motifs; the second learns compositional patterns over these features. Global max pooling provides position-invariant representations, and fully connected layers produce multi-label predictions.

#### 2.2.1 Embedding

Each amino acid is represented as a 34-dimensional vector combining one-hot encoding (20 dimensions) with 14 physicochemical properties: charge, hydrophobicity, polarity, aromaticity, hydrogen bond donor/acceptor capacity, molecular volume, special residue indicators (glycine, proline, cysteine), size category, and secondary structure propensities (helix, sheet, turn). This enhanced encoding provides the model with biochemically relevant features while remaining interpretable. Sequences are placed at random with periodic boundaries during training and zero-padded to a maximum length of 50 AA.

#### 2.2.2 Feature Extraction

The input representation is first projected to a 64-dimensional embedding space via a linear layer. Feature extraction then proceeds through two hierarchical convolutional blocks:

##### Multi-scale convolution block 1

Four parallel 1D convolutional branches with kernel sizes *k* ∈ {3, 5, 7, 9} and 128 filters each, followed by batch normalization and ReLU activation. These kernel sizes capture local sequence windows ranging from 3 to 9 residues, aligning with the 4–8 residue functional motifs identified in our dataset analysis. Longer kernel sizes can sense compositional improbabilities of decoy samples. The outputs are concatenated along the channel dimension, yielding a 512-dimensional feature map.

##### Multi-scale convolution block 2

A second set of parallel convolutions with identical configuration operates on the concatenated features from block 1. This hierarchical structure enables the network to learn compositional patterns over the motifs detected in the first layer. Each branch applies global max pooling over the sequence dimension, extracting the strongest activation for each filter regardless of position—appropriate for peptides where functional motifs can occur at varying locations.

#### 2.2.3 Classification Head

The pooled features (512 dimensions) pass through two fully connected layers (1024 and 512 units) with ReLU activation and dropout (*p* = 0.1), followed by a final linear layer mapping to 49 output labels. Sigmoid activation produces independent probabilities for each therapeutic function, enabling multi-label prediction. At inference time, labels with predicted probability ≥0.5 are assigned to the input sequence.

A smaller version of this architecture with approximately 1M parameters was used as the baseline configuration, whereas the fine-tuned model (Fig. 4) contains approximately 3M parameters per ensemble member and 15M parameters in total.

### 2.3 Loss Function

Multi-label classification with severe class imbalance requires careful loss design. We employ weighted Binary Cross-Entropy (BCE) loss, computed independently for each of the *C* = 49 labels:

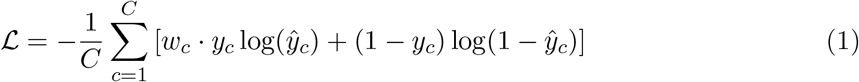

where *y*_*c*_ ∈ {0, 1}is the ground truth for label *c, ŷ*_*c*_ = *σ*(*z*_*c*_) is the predicted probability after sigmoid activation, and *w*_*c*_ is a class-specific weight that upweights positive samples for rare classes.

The class weights *w*_*c*_ are derived from the positive-to-negative ratio for each label. We investigate a weighting scheme of increasing strength. *No weighting* (*w*_*c*_ = 1) treats all samples equally, optimizing for overall accuracy but potentially neglecting rare classes. *Very low weighting* applies a dampened correction 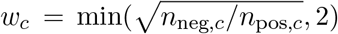, providing minimal emphasis on rare classes. *Low weighting* uses the same square-root formula with a cap of 5, providing moderate emphasis while preventing extreme weights. *Medium weighting* increases the cap to 10, allowing stronger correction for highly imbalanced labels. *Strong weighting* increases the cap to 15, more aggressively prioritizing rare class recall at potential cost to common class precision. All weighted schemes use square-root dampening, reflecting the empirical observation that full inverse-frequency weighting often overcorrects, degrading overall performance.

### 2.4 Training Details

We optimize using AdamW with an initial learning rate of 10^*−*3^ and weight decay of 10^*−*5^. Training proceeds with mini-batches of 64 samples, randomly shuffled each epoch. To prevent overfitting, we employ early stopping with a patience of 10 epochs, monitoring validation Micro F1 (defined in Section 3.2) as the stopping criterion. The learning rate is reduced by a factor of 0.5 when validation performance plateaus for 5 consecutive epochs (ReduceLROnPlateau scheduler). Training runs for a maximum of 60 epochs, though early stopping typically terminates training between epochs 30–50. The dataset is split into training (80%), validation (10%), and test (10%) sets with stratified sampling to preserve label distributions.

For the fine-tuned model, we additionally employ random start position augmentation: peptide sequences are randomly positioned within the input window rather than always left-aligned, encouraging the model to learn position-invariant features. We used periodic boundaries upon placement which ensures that every sequence can be placed at any position for any sequence length.

#### Implementation

Our implementation uses PyTorch [Paszke et al., 2019] for model development and training. Data preprocessing and analysis employ NumPy [Harris et al., 2020] and pandas [pandas development team, 2020], while evaluation metrics are computed using scikit-learn [Pedregosa et al., 2011]. All experiments were conducted on NVIDIA RTX 5070 GPUs.

## 3 Experiments

### 3.1 Experimental Setup

Our experimental methodology proceeds in three phases. In the *exploratory phase*, we conducted unstructured experiments across different architectures, encodings, and hyperparameters to identify a stable baseline configuration. This phase revealed that CNN-based architectures consistently outperformed transformers and that enhanced encodings provided marginal benefits over basic one-hot representations (please see Appendix Section A.1 for a comparison of different architectures).

Building on these findings, we proceeded to the *systematic evaluation phase*, where we isolate individual factors while holding others constant. Starting from a reference configuration (CNN architecture, enhanced encoding, medium class weighting, combined negative sampling), we vary one dimension at a time to quantify its effect on model performance. This controlled approach enables principled conclusions about each design choice.

Finally, in the *fine-tuning phase*, we combine the insights from systematic evaluation and iteratively evaluated different combinations of parameters to arrive at an optimized model with the best-performing settings identified for each dimension.

The systematic evaluation examines four dimensions, each addressing a specific research question:

#### Architecture comparison

We evaluate whether complex architectures provide benefits over simple CNNs for therapeutic peptide classification. Beyond our proposed two-layer CNN, we test CNN-Dilated (adding dilated convolutions to capture longer-range patterns), CNN-Attention (incorporating self-attention after convolution), ResNet (adding residual connections), and a Transformer encoder following the design principles of TPpred-LE. If local sequence motifs are indeed sufficient for this task, simpler architectures should perform comparably to or better than attention-based models.

#### Input encoding

We assess whether incorporating domain knowledge through physicochemical properties improves over pure sequence information. We compare basic one-hot encoding (20 dimensions), our enhanced encoding with physicochemical properties (34 dimensions), and learned embeddings of varying sizes (64 and 128 dimensions). This comparison reveals whether the model benefits from explicit biochemical features or can learn equivalent representations from data.

#### Class weighting

Given the severe class imbalance in therapeutic peptide data, we investigate how different weighting schemes affect the trade-off between overall accuracy (Micro F1) and balanced per-class performance (Macro F1). Understanding this trade-off is crucial for practical applications where rare therapeutic functions may be of particular interest.

#### Dataset composition

We validate our negative sampling strategy through ablation studies. By training models with individual sampling methods and measuring performance when each method is removed from the combined baseline, we quantify the contribution of each decoy generation approach. We also examine how the positive-to-negative ratio affects model behavior.

### 3.2 Evaluation Metrics

We evaluate model performance using precision, recall, F1 score, and false positive rate. Given true positives (*TP*), false positives (*FP*), false negatives (*FN*), and true negatives (*TN*), precision measures the fraction of positive predictions that are correct, while recall (also called the true positive rate) measures the fraction of actual positives that are identified:

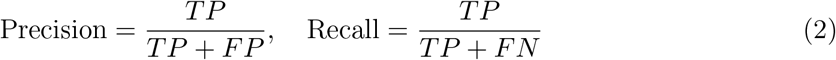

The F1 score is the harmonic mean of precision and recall, balancing both metrics:

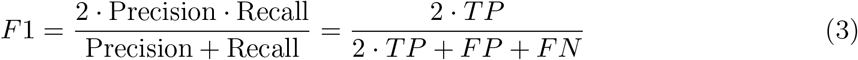

Complementing recall, the false positive rate (FPR) measures the fraction of true negatives that are incorrectly predicted as positive:

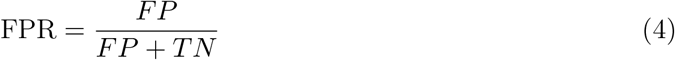

Whereas precision is sensitive to the prevalence of positives, FPR is independent of class balance and is therefore informative when evaluating models on label sets with widely varying support, or when the cost of false alarms must be controlled directly.

For multi-label evaluation, we report two complementary aggregation strategies; Micro and Macro F1. The tension between these metrics, where improving one often degrades the other, motivates our investigation of class weighting strategies. Micro F1 computes precision and recall by summing *TP, FP*, and *FN* across all labels before computing the score, effectively weighting each label by its frequency. This metric reflects overall prediction quality and is dominated by common classes. Macro F1 computes F1 independently for each label and averages the results, giving equal weight to rare and common classes alike.

### 3.3 External Validation Protocol

To assess generalization beyond our training distribution, we evaluate on the independent test set from TPpred-LE. This external benchmark uses different data sources and annotation protocols, providing a rigorous test of model transferability.

A key challenge is label reconciliation: our model predicts 48 therapeutic function labels plus one global therapeutic flag, while the TPpred-LE benchmark defines 15 categories, 12 of which admit an unambiguous semantic mapping. We established a mapping between the 12 overlapping label categories based on semantic equivalence (e.g., our “Antibacterial” maps to TPpred-LE’s “ABP”). The table can be found in the Appendix in Section A.4. For evaluation, we extract only the predictions corresponding to these 12 shared labels and compute metrics on this subset. This setup is conservative - our model solves the more challenging task of predicting 49 labels, yet we evaluate on only 12. Labels unique to either dataset are excluded from comparison.

This external validation complements our held-out test set evaluation by measuring performance on data collected and annotated independently, revealing whether our model has learned generalizable features of therapeutic peptide function rather than dataset-specific artifacts.

## 4 Results and Discussion

We present a comprehensive evaluation of our CNN-based approach for multi-label therapeutic peptide classification. Detailed results of the model comparison, systematic hyperparameter tuning, and finetuning are shown in the Appendix in Section A.2. Here, we will present a summary of the benchmark comparisons and present detailed results of the final model.

The model comparison revealed that CNN architectures systematically outperform transformer and CNN-attention models in terms of F1 scores and parameter efficiency. 2-Layer CNN-architectures found to consistently provide good performance even for small models with a strong response to increasing number of parameters. Adding dilated layers or residual layers did not measurably improve the performance and were thus dropped for simplicity. Optimal parameters were found through a systematic hyperparameter search and subsequent finetuning as described in Section 3. The architecture of the final model which we refer to as *TheraPep-AI* is presented in Fig. 4.

We evaluated *TheraPep-AI* on the TheraPepDB dataset; the results are shown in Table 1. The model achieved 78.9% Micro F1 and 54.4% Macro F1 across 48 therapeutic categories and a 2.1% FPR on the augmented negative data. On the combined positives and negatives the performance is 78.9% Micro F1 and 54.4% Macro F1. To the best of our knowledge, this represents the largest reported evaluation of therapeutic peptide classification to date, both in terms of the number of functional categories and training set size. The 24.5% difference between Macro and Micro F1 highlights that highly populated classes are predicted more reliably than data with low population. Neuropeptides stand out as the most reliably predicted category with 84% F1 which we attribute to the importance of sequence motifs, their compositional dissimilarity to the other classes (see. Fig. 3), and their abundance in the dataset. As expected, very low populated classes have a significantly lower performance. The lowest performance is observed for Antiinflammatory peptides (<300 sequences) with around 7.1 %.

**Table 1:** Comparison with State-of-the-Art Methods. F1 scores on positive therapeutic peptides (15 major categories (bold face) and 11 of 32 subcategories with larger support) and FPRs on synthetic negatives. All were trained, validated, and tested on TheraPepDB with identical splits (80% train, 10% val, 10% test).

| Label | Test Support<sup>†</sup> | TPpred-LE<br>(retrained) | PrMFTP<br>(retrained) | Ours<br>(Fine-tuned) |
| --- | --- | --- | --- | --- |
| *Positive Samples – F1 scores (%)* |  |  |  |  |
| **Antibacterial** | 2,020 | 76.3 | 57.7 | **78.7** |
| **Antimicrobial** | 3,863 | 88.3 | 14.2 | **90.0** |
| **Neuropeptide** | 821 | 78.8 | 63.6 | **85.3** |
| **Antifungal** | 706 | 53.0 | 10.7 | **59.4** |
| **Antiviral** | 651 | 63.8 | 49.6 | **70.8** |
| **Anticancer** | 608 | 52.0 | 18.9 | **56.3** |
| **Antihypertensive** | 346 | 51.0 | 21.1 | **65.4** |
| **Metabolic regulatory** | 268 | 49.7 | 29.7 | **61.7** |
| **Cell communication** | 213 | **52.0** | 17.8 | 51.9 |
| **Drug delivery** | 143 | 54.2 | 47.9 | **55.1** |
| **Antioxidant** | 140 | 16.2 | 0.0 | **27.6** |
| **Antiparasitic** | 86 | 45.0 | 33.3 | **49.2** |
| **Immunoactive** | 77 | 41.4 | 39.6 | **49.2** |
| **Growth Regeneration** | 13 | **66.7** | 63.2 | **66.7** |
| **Thrombolytic** | 19 | 28.3 | 0.0 | **41.4** |
| AntiHIV | 231 | 69.3 | 63.4 | **76.4** |
| Glucose metabolism | 148 | 49.4 | 40.0 | **64.2** |
| Cell-penetrating | 77 | 62.9 | 56.1 | **66.7** |
| AntiHCV | 60 | 86.8 | 86.8 | **87.9** |
| Tumor homing | 58 | **54.5** | 38.9 | 41.0 |
| AntiBreastcancer | 40 | 63.2 | 61.0 | **63.9** |
| Anti-angiogenesis | 31 | 35.9 | 27.8 | **50.0** |
| AntiLungcancer | 30 | 51.0 | 57.1 | **59.3** |
| Quorum sensing | 27 | **63.6** | 41.2 | 61.5 |
| AntiColoncancer | 27 | 65.5 | **71.4** | 66.7 |
| Antiinflammatory | 21 | 0.0 | 0.0 | **7.1** |
| **Micro F1 (positives)** | 5,566 | 77.8 | 50.3 | **79.9** |
| **Macro F1 (positives)** | 5,566 | 47.7 | 38.7 | **54.6** |
| *Negative Samples* |  |  |  |  |
| **Overall FPR** | 10,971 | 13.9 | 1.0 | **2.1** |
| *Full Test Set* |  |  |  |  |
| **Micro F1 (full)** | 16,537 | 71.0 | 49.3 | **78.9** |
| **Macro F1 (full)** | 16,537 | 44.6 | 38.5 | **54.4** |

<sup>†</sup> Per-label test support is the number of test peptides annotated with that label. As peptides are multi-label, these counts overlap and sum to more than the 5,566 positive samples. The summary rows instead report sample counts.

Some performance degradation likely originates from data uncertainty: many functions are set to negative (0) if no tests have been found for them. Across all categories, this is an assumption that holds on average but not for every subcategory. Many therapeutic peptides started as derivatives of extracted natural peptides of fungi or other organisms. An anti cancer drug might have started from an anti microbial drug and been modified to increase specificity, lifetime, or other properties. Experiments for the original category which was of no or low monetary value (for example, antimicrobial function) might have been skipped and consequently those are reported as negatives. Since the most prevalent classes are also the ones with lowest monetary value, they likely limit performance of the rare but important classes. We consider this data uncertainty limitation that we do not address in the present work but where differentiating labels between validated negatives and unknown targets could help to improve performance. We used a 2:1 negative to positive data ratio in training which ensures that the model defaults to negative data in uncertain cases due to statistical likelihood. Due to the statistical features of the augmented data, the model has to pick up on biologically relevant motifs instead of just focusing on compositional features alone. In later sections we analyze this in more detail.

We compare our model to the foundational work of TPpred-LE [Lv et al., 2023b] and PrMFTP [Yan et al., 2022]. The first is a transformer encoder-decoder model whereas the second is a multi-scale CNN + bi-directional LSTM + multi-head self-attention mode. To establish a fair comparison with prior methods, we trained TPpred-LE and PrMFTP [Yan et al., 2022] on our TheraPepDB dataset using identical train/validation/test splits, then evaluated all models on our held-out test set across all 49 therapeutic categories. To enable a comparison between all categories, we extended the output layers to match our labels. This allows for a direct comparison between the different architectures while neglecting the differences resulting from the drastically larger positive datasets and contrast provided by the augmented negative datasets. Table 1 shows the results for TPpred-LE and PrMFTP. TPpred-LE achieves almost the same performance as *TheraPep-AI* on positive data but struggles distinguishing the positive samples from negative ones (13.9% FPR opposed to 2.1%). PrMFTP performs significantly worse on the positive data but achieves an excellent FPR of 1.0%. Overall our CNN architecture significantly outperforms previous models when trained on the same data.

On positive samples, our model achieves 79.9% Micro F1 and 54.6% Macro F1, compared to 77.8%/47.7% for TPpred-LE and 50.3%/38.7% for PrMFTP. The improvement is particularly notable for rare categories such as Antihypertensive, Anti-angiogenesis, and Glucose metabolism, where our class-balanced training strategy appears beneficial.

On the synthetic negatives, the PrMFTP model performs best, with an FPR of 1.0%, whereas our model obtains an FPR of 2.1%. The transformer architecture of TPpred-LE performs substantially worse than both PrMFTP and our model, with an FPR of 13.9%. Since the synthetic data are generated based on compositional and neighboring transition statistics, this finding indicates that these features play a more central role in the transformer architecture than in the other models. We hypothesize that CNNs benefit from *translational invariance*— detecting functional motifs regardless of their position in the sequence—whereas transformers with positional encodings may learn position-specific patterns that fail to generalize. We explore this hypothesis further in Section A.1.

The low FPR of CNN-based models suggests that training with diverse synthetic negatives teaches these architectures to recognize what makes a peptide *therapeutic* rather than merely *peptide-like*. This distinction is crucial for practical applications where the model must screen large libraries of candidate sequences. On the full test set (including synthetic negatives), we observe that FPRs significantly impact overall performance. Our model maintains 78.9% Micro F1, TPpred-LE achieves 71.0 % Micro F1 and PrMFTP obtains 49.3 %. We note that this comparison is conducted on our dataset; different training data distributions may favor different architectural choices and will be analyzed below.

### 4.1 External Benchmark Validation

Performance on randomly sampled test data from the same distribution as training data may overestimate a model’s true generalization capability. Test sequences that share similar features with training data—due to common sources, similar experimental conditions, or database overlap—can inflate accuracy metrics without revealing how the model performs on genuinely novel sequences. To assess generalization more rigorously, we evaluate on an external benchmark derived from an independent data source.

We use the TPpred-LE benchmark [Lv et al., 2023a] as our external test set (1,024 sequences, 12 therapeutic categories). Initial analysis revealed that TheraPepDB and the TPpred-LE benchmark share some overlapping sequences. To ensure a fair evaluation, we removed all sequences from our training data that appear in the TPpred-LE benchmark, then retrained both our CNN and the TPpred-LE architecture on this cleaned dataset. This guarantees 0% sequence overlap between training and evaluation, providing a stringent test of generalization.

Table 2 compares three models: TPpred-LE trained on its original dataset (12 labels), TPpred-LE retrained on our cleaned TheraPepDB (49 labels), and our CNN trained on the same cleaned data. All models were evaluated on the test set of the 12 label dataset and shown for comparison. The 49 labels were mapped to the 12 labels.

**Table 2:** TPpred-LE Benchmark Results. Comparison on the TPpred-LE external benchmark (1,024 sequences, 12 labels). All models evaluated have 0% training data overlap with this benchmark.

| Label | Function | Support | TPpred-LE<br>(internal) | TPpred-LE<br>(external) | **Ours**<br>(external) |
| --- | --- | --- | --- | --- | --- |
| *Training dataset* |  |  | TPpred-LE | TheraPepDB | TheraPepDB |
| *F1 scores (%)* |  |  |  |  |  |
| AMP | Antimicrobial | 500 | 74.6 | 65.6 | **73.2** |
| ABP | Antibacterial | 150 | 40.8 | 25.6 | **44.8** |
| AIP | Antiinflammatory | 149 | **63.1** | 25.4 | 38.5 |
| AVP | Antiviral | 139 | 57.5 | 23.6 | **56.2** |
| ACP | Anticancer | 83 | 39.0 | 15.0 | **42.2** |
| AFP | Antifungal | 73 | 20.2 | 13.3 | **36.9** |
| DDV | Drug delivery | 59 | 44.0 | 10.5 | **46.0** |
| CPP | Cell-penetrating | 50 | **54.2** | 9.3 | 44.0 |
| APP | Antiparasitic | 14 | 0.0 | 2.8 | **17.4** |
| AAP | Anti-angiogenic | 13 | 28.6 | 2.4 | **27.3** |
| AHTP | Antihypertensive | 6 | 0.0 | 1.2 | **14.3** |
| QSP | Quorum sensing | 6 | **50.0** | 1.1 | 22.2 |
| **Micro F1** |  | 1024 | **57.9** | 18.8 | 55.3 |
| **Macro F1** |  | 1024 | 38.1 | 16.3 | **38.6** |

<sup>†</sup> Per-label test support is the number of test peptides annotated with that label. As peptides are multi-label, these counts overlap and sum to more than the 1,024 test peptides. The summary rows instead report sample counts.
TPpred-LE (original): trained on the TPpred-LE dataset (12 labels). TPpred-LE (retrained) and Ours: trained on TheraPepDB (49 labels), evaluated on the 12 shared labels. All training sets have 0% sequence overlap with this benchmark.

The results reveal striking differences in generalization capability. TPpred-LE trained on its native 12-label dataset achieves the best Micro F1 (57.9%), demonstrating the advantage of domain-specific training where test data closely matches the training distribution. However, when both architectures are trained on TheraPepDB and evaluated on this external benchmark, our CNN achieves 55.3% Micro F1 while the TPpred-LE transformer drops to 18.8%—a 3 × difference in generalization performance.

This performance gap is particularly notable because both models were trained on identical data with 0% overlap with the benchmark. The CNN maintains near-parity with the domain-specific TPpred-LE (55.3% vs 57.9%), while the transformer architecture fails to transfer its learned representations to the external test set. On rare classes where TPpred-LE (origi-nal) achieves 0% F1 (Antiparasitic, Antihypertensive), our model achieves non-zero predictions (17.4%, 14.3%), suggesting better handling of low-frequency categories.

These results provide evidence that CNN architectures generalize better than transformers for therapeutic peptide classification. The external benchmark represents a stringent test: sequences from a different data source, with different collection criteria and potentially different sequence characteristics. The transformer’s dramatic performance drop (from competitive on internal test sets to 18.8% on external data) suggests it learns dataset-specific patterns rather than generalizable sequence-function relationships.

We attribute this difference to translational invariance. The CNN’s convolutional filters detect motifs regardless of their position. Position-invariant features transfer naturally to new datasets where the same motifs may appear at different locations. The transformer’s positional encodings, while powerful for tasks requiring position-aware reasoning, may be a liability when positions carry no biological meaning—as is often the case for short therapeutic peptides where functional motifs can occur anywhere in the sequence.

### 4.2 False Positive Analysis: Why Negative Sampling Matters

A critical consideration for therapeutic peptide prediction and a main consideration of this work is the FPR – how often the model incorrectly predicts therapeutic activity for non-therapeutic sequences. Our model achieves a low FPR of 2.1%, which we attribute to our diverse negative sampling strategy during training. We analyze the impact of different negative sampling configurations below.

As a baseline, we trained the same model on the positive data alone and compared to the model that was trained on the positive and negative data. When evaluated on positive samples only, the positive-only model achieves 78.8/54.5% micro/macro F1 on the 48 therapeutic labels (leaving out the label determining therapeutic function which would always be true) whereas the model trained on both positive and negative data achieves 75.8/53.9% micro/macro F1. However, when evaluating the positive-only model on negative data - counting a sequence as a correct rejection when no therapeutic function label exceeds the threshold (the flag denoting therapeutic function is excluded, as the positive-only model trivially predicts it for every input) - we obtain an 88% FPR, compared to 1.5% for the model trained on both positives and negatives under the same criterion. Thus a −3.0/-0.6% drop in positive-classification F1 is traded for a strong reduction in false positives on the synthetic negatives.

Figure 5a reveals the effect of training on only a single negative sampling method. We trained the finetuned model multiple times by including only a single type of augmented negative data (uniform, global Markov, position Markov, class frequency). Training exclusively on Uniform random negatives leads to catastrophic generalization failure: FPR reaches 26–31% on Markov, Position-Markov, and Class-Frequency negatives. This occurs because uniform random sequences lack peptide-like compositional structure—they are “too easy” to reject, and the model fails to learn features that distinguish realistic non-therapeutic sequences. In contrast, models trained on any single realistic method (Markov, Position-Markov, or Class-Frequency) achieve moderate cross-type generalization (2–6% FPR). We note that, although the dataset contains many short peptides (<10 amino acids), compositional features can still be statistically informative because each residue position is sampled from a 20-letter amino acid alphabet.

**Figure 5:**
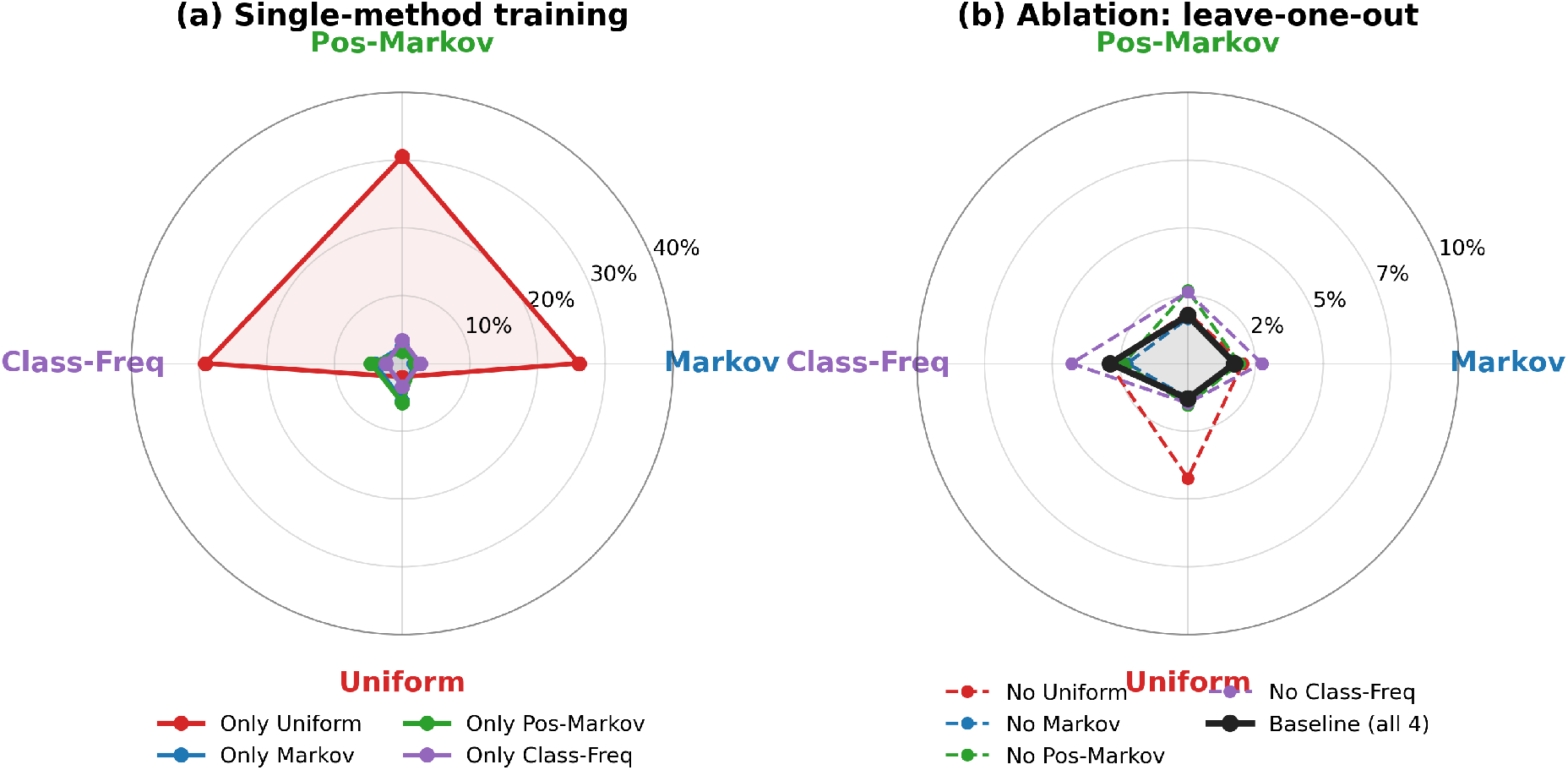
False-positive-rate (FPR) analysis of negative-sampling strategies. (a) Single-method training: training on Uniform negatives alone (red) causes catastrophic failure on the three structured negative types (26–31% FPR), whereas the other single methods generalize moderately (2–6% FPR). (b) Leave-one-out ablation: FPR when each negative-sampling method is excluded from training. Removing a method raises FPR most on the negative type it targets (e.g. omitting Uniform or Class-Frequency lifts their respective FPRs to ∼4%), while the baseline trained on all four types (black) achieves consistently low FPR (≤2.9%) across all types.

Figure 5b shows the FPR obtained when each negative sampling method is individually excluded from training. The baseline model, trained on negatives from all four methods, maintains consistently low FPR across all negative types (1–3%). In contrast, omitting a given method tends to increase the FPR on negatives generated by that same method. The effect is largest for Uniform: excluding Uniform negatives raises the FPR on Uniform test samples from 1.3% to 4.2%. Excluding Class-Frequency or Position-Markov negatives produces smaller, method-specific increases (Class-Frequency 2.9% to 4.3%, Position-Markov 1.8% to 2.7%), whereas excluding global Markov has little effect (1.7% to 1.9%). These results suggest that the position-, class-frequency-, and global-Markov-based negatives occupy more similar regions of sequence space, whereas Uniform negatives provide a more distinct background distribution. Including Uniform negatives may therefore help the model define a broader negative sequence space and prevent random amino acid sequences from being ranked as positives.

These results validate our diverse negative sampling strategy: combining multiple methods yields a model that robustly rejects non-therapeutic sequences regardless of their compositional sophistication. The gradient of difficulty—from easy (Uniform) to hard (Class-Frequency)— forces the model to learn discriminative features at multiple levels of abstraction, preventing overfitting to superficial patterns.

The importance of negative sampling extends beyond classification metrics. In practical drug discovery, the cost of false positives (synthesizing and testing inactive peptides) far exceeds the cost of false negatives (missing some candidates). Our analysis shows that naive negative sampling creates “shortcut” learning—the model distinguishes therapeutic peptides from random sequences using simple compositional features (amino acid frequencies) rather than functional motifs. When presented with realistic non-therapeutic sequences that share peptide-like composition, such models fail catastrophically. The diverse negative sampling strategy forces the model to learn deeper features that distinguish therapeutic function from mere peptide structure.

### 4.3 Error Analysis: Label Confusion Patterns

To understand systematic prediction errors, we analyze multi-label confusion patterns on the held-out test set. For readability, Figure 6 displays the 22 labels with sufficient test support (≥ 50 samples), ordered hierarchically.

**Figure 6:**
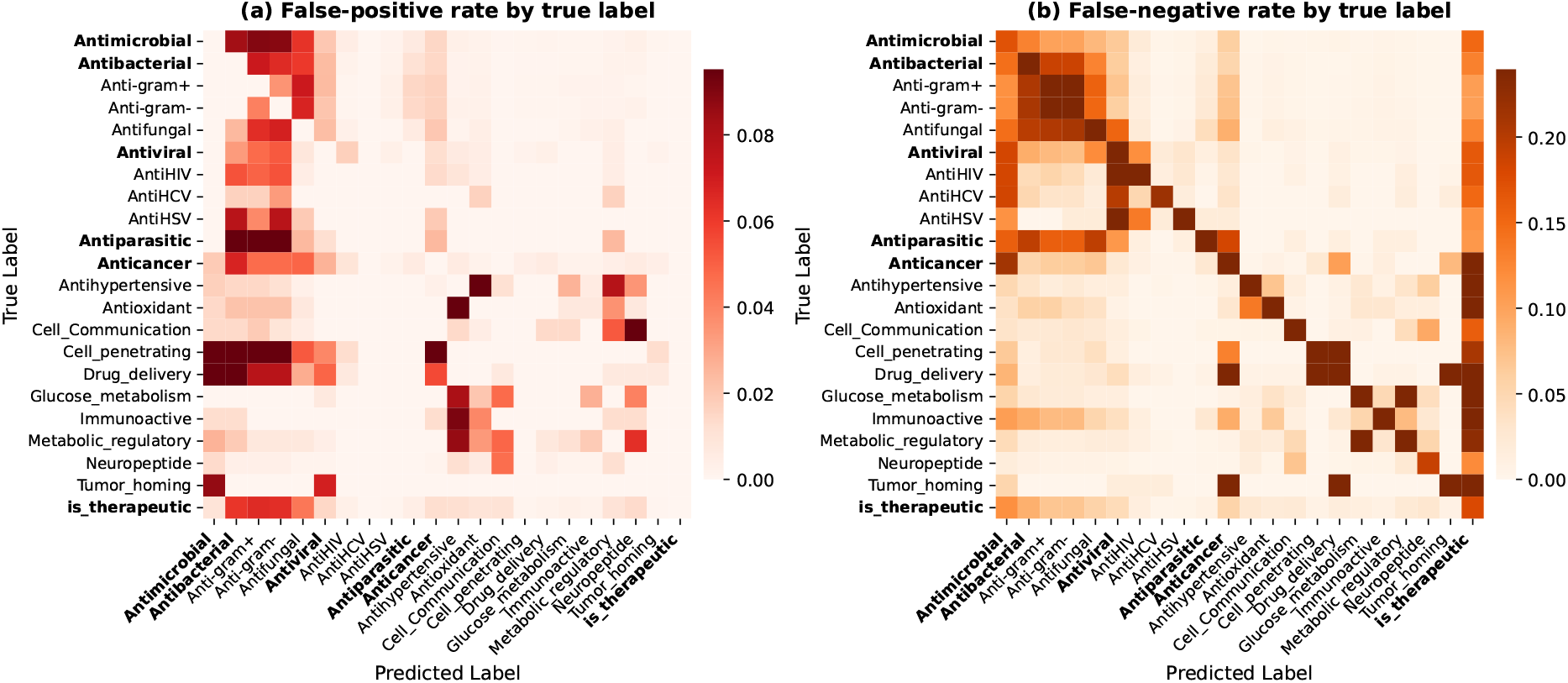
Multi-Label Confusion Analysis. (a) False-positive patterns: for samples with true label *i* (row), the rate at which an absent label *j* (column) is incorrectly predicted. (b) False-negative patterns: for samples with true label *i* (row), the rate at which a co-occurring true label *j* is missed. Both are normalized by the true-label support. Most confusion occurs within biological families (antimicrobial, anticancer), reflecting genuine functional overlap.

#### Confusion concentrates within biological families

The strongest false-positive signals occur between functionally related labels rather than across unrelated families. Within the antimicrobial group, for instance, antiparasitic peptides are predicted as antibacterial or Antigram+ at ∼17%, and cell-penetrating peptides trigger antimicrobial/antibacterial predictions at ∼18%. Because the label set is hierarchically propagated (a subtype always implies its parent), parent labels such as Anticancer are never false positives for their subtypes; instead, the parent direction appears in the false-negative panel, where specific labels are predicted while the broader umbrella is dropped (e.g. AntiHSV misses its Antiviral parent at 31%, and Glucose-metabolism misses Metabolic-regulatory at 38%).

#### Multi-function peptides drive false negatives

Peptides annotated with several cooccurring functions are the hardest to fully recover: tumor-homing peptides miss their co-labels Drug-delivery and Anticancer at ∼72%, and the overarching is_therapeutic flag is the single most frequently missed label. This reflects the difficulty of simultaneously predicting all members of a large co-occurring label set.

#### Major families remain well separated

Direct confusion between the two dominant families—antimicrobial and anticancer—is low (∼1.5–1.9%), indicating the model distinguishes their fundamentally different mechanisms of action rather than relying on superficial sequence similarity. Some functionally adjacent categories do show higher cross-prediction (e.g. antioxidant versus antihypertensive, ∼12–14%), consistent with genuine biological overlap.

#### Rare classes remain challenging

The full 49-label matrices show that labels with very few test examples are recovered poorly: AntiTubercular (*n* = 2) and Osteogenic (*n* = 3) reach 0% recall, and other low-support categories remain weak (e.g. Antimalarial 17%, *n* = 6; Antileishmania 20%, *n* = 5). This motivates future work on few-shot or transfer learning for rare therapeutic functions.

### 4.4 Implicit Detection of Structural and Electrostatic Effects

Beyond sequence motifs, the model captures biophysical determinants of therapeutic function. Many antimicrobial peptides act through the conserved **WxxWxxW** pattern [Tsai et al., 2022], in which three tryptophans spaced by three residues align on a single face of an *α*-helix and insert into the lipid bilayer, while flanking cationic residues (K, R) anchor the peptide at the negatively charged bacterial membrane. Disrupting either ingredient – the amphipathic helix or the net positive charge—is known to abolish activity.

Figure 7 compares five variants that all share the canonical W-spacing. Variants (a)–(c) preserve both the helical fold and the cationic surface and are correctly classified as therapeutic by both our model and TPpred-LE. Variant (d) replaces internal residues with glycines (KK W**G**N W**GG** WRL KK), which destabilize the *α*-helix and collapse the structure into a disordered loop, breaking the spatial alignment of the tryptophans. Variant (e) keeps the helix intact but substitutes lysines with the negatively charged aspartate and glutamate (KK W**KD** W**ES** WRL KK), neutralizing the cationic face required for membrane localization. We removed one of the sequences that was present in our training dataset from it and retrained. Our model assigns confidence below 0.3% to both (d) and (e), whereas TPpred-LE misclassifies them as therapeutic with 60% and 61% probability. For qualitative structural visualization, peptide structures in Figure 7 were predicted from amino-acid sequences using AlphaFold2; these structures were not used as model inputs or training features [Jumper et al., 2021].

**Figure 7:**
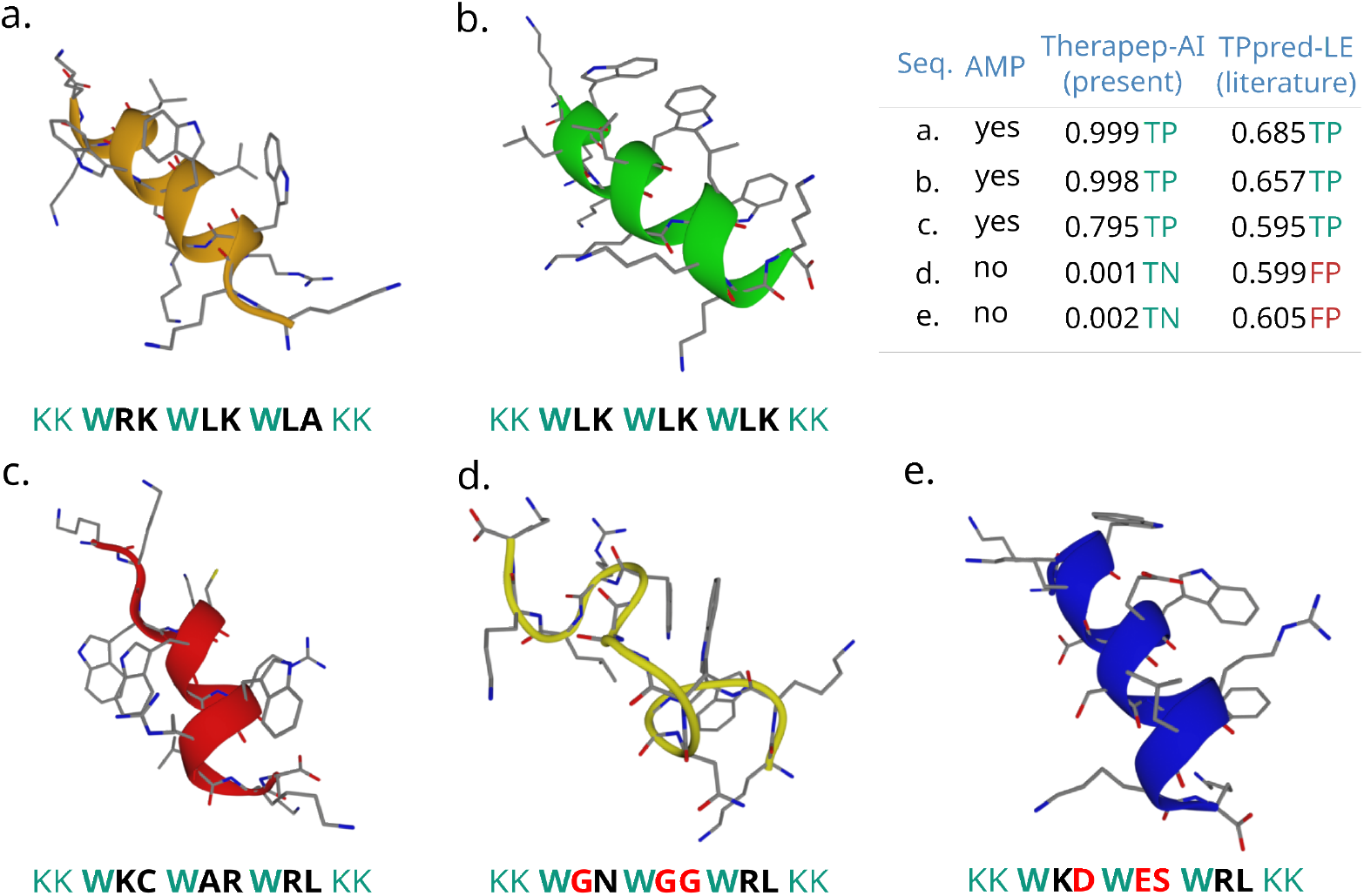
Implicit Detection of Structural and Electrostatic Effects. Predicted folds and classifier probabilities for five variants of an antimicrobial peptide sharing the WxxWxxW spacing. (a)–(c) preserve the amphipathic *α*-helix and the cationic flanking residues and are correctly predicted as therapeutic by both our model and TPpred-LE. (d) Glycine substitutions destabilize the helix into a disordered loop. (e) Negatively charged substitutions (D, E) preserve the helix but neutralize the cationic face required for membrane localization. Our model assigns confidence below 0.3% to both (d) and (e); TPpred-LE misclassifies both as therapeutic.

Because the network sees only the amino acid sequence and has no explicit structural input, this behavior indicates that it has internalized two distinct biophysical principles directly from sequence statistics: (i) the helix-breaking propensity of glycine, and (ii) the requirement for a net positive charge to localize at the membrane. We further hypothesize that the 2:1 negative-to-positive training ratio makes the model strict when deciding whether a sequence is therapeutic: the presence of the **WxxWxx W** pattern alone is not sufficient for a positive call, and the model additionally requires the surrounding context that, in the training distribution, co-occurs with a functional amphipathic helix. Together with the W-spacing detector, these implicit features let the model discriminate functional amphipathic helices from sequences that superficially preserve the W motif but lack the underlying biophysical context — a distinction that may be missed when superficial motif conservation dominates the prediction.

### 4.5 Model Interpretability

To understand what sequence patterns the CNN learns, we trained a variant with L1 regularization on neuron activations. This sparsity constraint encourages individual filters to specialize, making their function interpretable while maintaining competitive performance.

The L1-regularized model achieves 85–98% sparsity in convolutional layers (compared to ∼50% from ReLU alone), meaning only 2–15% of filters activate for any given peptide. This sparsity enables us to analyze what each filter has learned by correlating its activation patterns with therapeutic labels.

Figure 8 shows filter-label correlations for the second convolutional layer (Conv2, kernel size 7). Individual filters exhibit clear specialization: F69 correlates strongly with Lipid_metabolism (*r* = 0.40), F1 with Neuropeptide (*r* = 0.31), and F124 with AntiHIV (*r* = 0.30). Multiple filters (F65, F107, F18) specialize for Antimicrobial detection (*r* = 0.23–0.28), consistent with this being the most frequent label. Notably, filters show both positive correlations (detecting presence of a function) and negative correlations (detecting absence), suggesting the network learns discriminative features for distinguishing between therapeutic categories.

**Figure 8:**
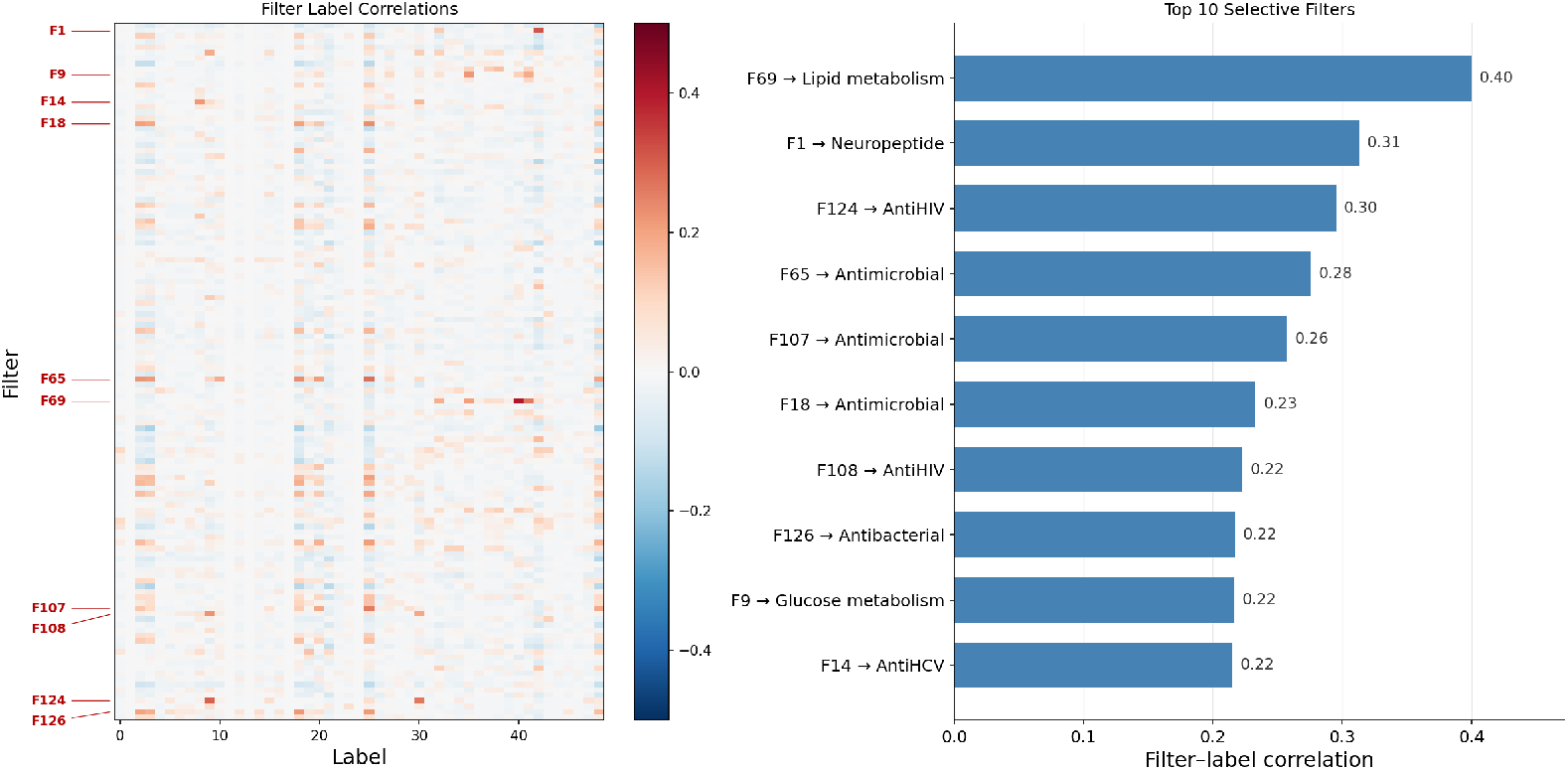
Filter-Label Correlations. (a) Pearson correlation between therapeutic labels and the activations of the 128 filters in one convolutional layer (Conv2, kernel size 7). Each row is a filter, each column a label. Filters show clear specialization: F69 for Lipid_metabolism, F1 for Neuropeptide, F124 for AntiHIV. Red indicates positive correlation (filter activates when label is present), blue indicates negative correlation. (b) The ten filters from this layer with the highest absolute filter–label correlation, each annotated with its most strongly associated therapeutic label.

#### 4.5.1 Learned Motif Detectors

The most striking finding is that specific filters learn to detect known biological motifs. Figure 9 shows activation patterns for five specialized filters across multiple therapeutic categories, with amino acid sequences overlaid on activation heatmaps.

**Figure 9:**
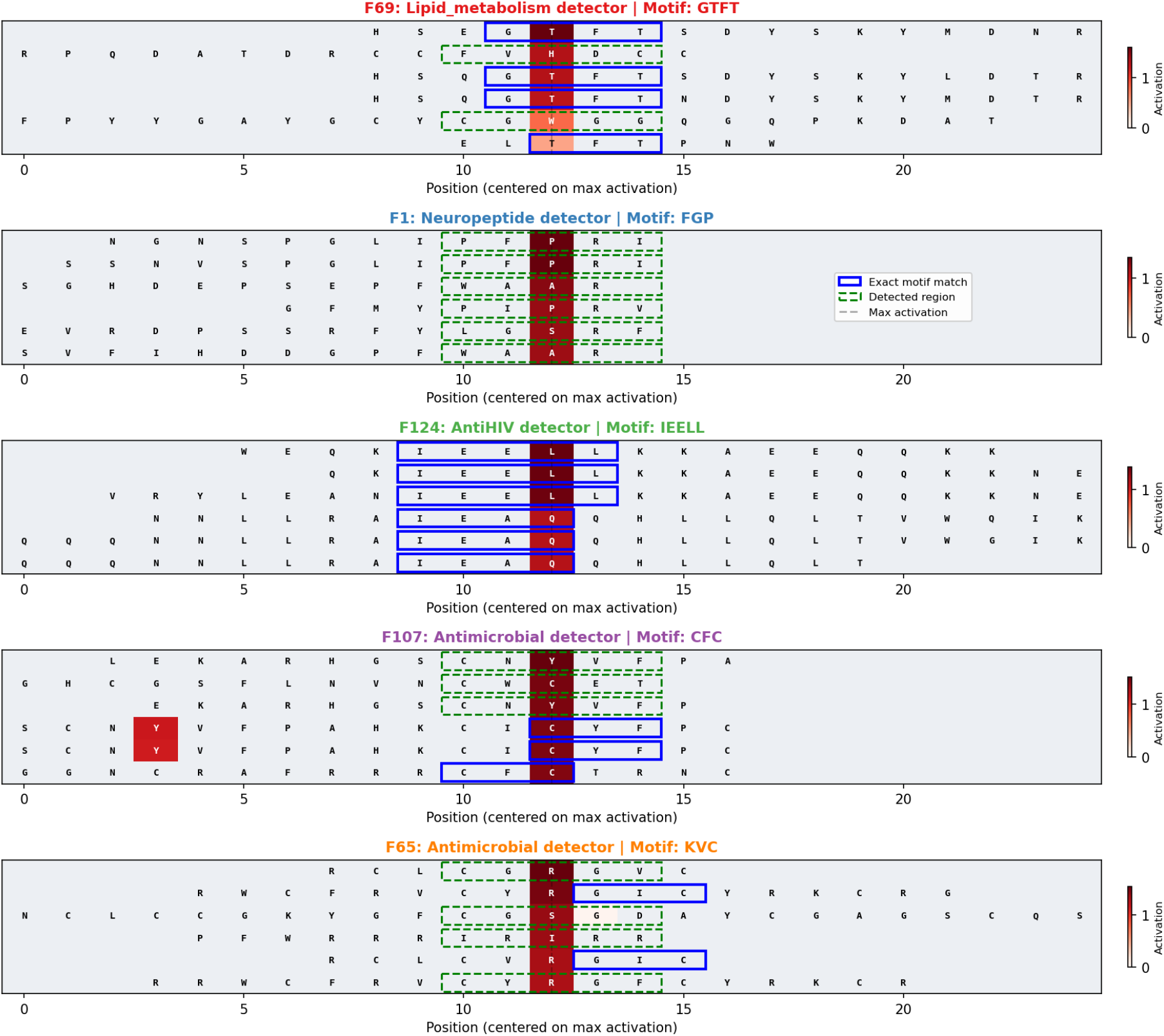
Learned Motif Detectors. Activation heatmaps for five specialized filters, each detecting distinct sequence motifs. Amino acid letters are overlaid on activation intensity (red = high activation). Blue boxes indicate exact motif matches; green dashed boxes show detected regions for motif variants. Each row shows a different peptide sequence, centered on maximum activation. F69 detects GTFT (glucagon-like signature), F1 detects FGP (neuropeptide motif), F124 detects IEELL (charged helix), F107 and F65 detect cysteine-rich patterns characteristic of antimicrobial peptides.

Each filter has learned to recognize biologically meaningful sequence patterns:

- **F69 (Lipid metabolism)**: Detects **GTFT/GTFTS**, the signature motif of glucagon-like peptides (GLP-1, GLP-2). Sequences containing this motif activate the filter ∼43× above baseline and show ∼100× enrichment for the Lipid_metabolism label.

##### F1 (Neuropeptide)

Detects aromatic–proline motifs of the **FGP** family (e.g. **WFGP, FGPR**), which activate the filter 18–20× above baseline. Its top-activating sequences are 84% Neuropeptide vs. 5% baseline.

- **F124 (AntiHIV)**: Detects **IEELL** and related charged coiled-coil motifs, likely involved in membrane interaction. Sequences containing it activate the filter *>*50× above baseline (59× label enrichment).
- **F107, F65 (Antimicrobial)**: Detect cysteine-rich, Cys–x–Cys patterns (e.g. **CFC** for F107) characteristic of defensins and other disulfide-stabilized antimicrobial peptides.

These motifs were learned without explicit supervision—the model discovered genuine sequence-function relationships from the training data alone.

#### 4.5.2 Interpretability-Performance Trade-off

The sparse model achieves 76% Micro F1 and 52% Macro F1, compared to 79% and 54% for the standard model—a modest 2pp reduction. This trade-off is acceptable for applications requiring model interpretability, such as understanding why a peptide is predicted to have specific therapeutic properties. The ability to identify which filters activate and what motifs they detect provides mechanistic insight into model predictions.

These results demonstrate that our CNN learns biologically meaningful features rather than spurious correlations. The emergence of motif detectors for enkephalin—without any explicit supervision for this pattern—suggests the model discovers genuine sequence-function relationships from the training data.

The interpretability analysis addresses a common concern about deep learning in biology: that models learn “black box” features with no biological meaning. Our sparse CNN variant demonstrates that interpretability and performance need not be mutually exclusive. The 2pp performance reduction is a modest cost for the ability to explain predictions in terms of known biological motifs. This has practical implications: when the model predicts a peptide as antimicrobial, we can examine which filters activated and whether they detected cysteine frameworks, amphipathic patterns, or other known antimicrobial features. Such explanations build trust in model predictions and can guide experimental validation. The discovery of the GTFT motif as a strong predictor of lipid metabolism function—recapitulating decades of glucagon research— validates that the model has learned genuine biology rather than dataset artifacts.

### 4.6 Summary

Our results demonstrate that CNN-based classification achieves strong performance on multilabel therapeutic peptide prediction, with 79.9/54.6% Micro/Macro F1 across 48 therapeutic function labels plus one global therapeutic flag. Three key design choices contribute to this performance: (1) diverse negative sampling that teaches the model to distinguish therapeutic function from peptide-like composition; (2) class-balanced loss weighting that prevents the model from ignoring rare categories; and (3) multi-scale convolutional filters that capture motifs at the relevant length scales (3–9 residues). The model’s learned features correspond to known biological motifs, providing confidence that predictions reflect genuine sequence-function relationships rather than spurious correlations.

#### Limitations

While our synthetic negative sampling enables FPR estimation, performance on real biological non-therapeutic sequences remains untested without experimental validation. The model struggles with rare therapeutic categories that have fewer than 10 training examples, achieving 0% recall on the rarest labels. Additionally, performance on novel therapeutic categories not represented in TheraPepDB cannot be assessed—the model can only predict functions it has seen during training. Finally, any candidate peptides identified through *in silico* screening should undergo wet-lab validation before therapeutic development, as computational predictions alone cannot guarantee biological activity.

## 5 Conclusion

We presented TheraPep-AI, a lightweight CNN for multi-label therapeutic-peptide function prediction that matches or exceeds transformer-based methods at a fraction of their parameter count. Through systematic experiments we established four design principles:

1. Simple multi-scale CNN architectures outperform attention- and transformer-based models on this task;
2. Diverse negative sampling (uniform, Markov, position-Markov, and class-frequency) is essential for generalization, reducing the false-positive rate on synthetic negatives from ∼88% (positive-only training) to 2.1%;
3. Square-root class weighting with a cap of 2 gives the best Micro/Macro F1 balance;
4. Physicochemically enhanced encodings are competitive with learned embeddings, indicating that domain knowledge remains valuable.

On the TheraPep-DB test set the fine-tuned model reaches 79.9% Micro and 54.6% Macro F1 while maintaining a 2.1% false-positive rate on held-out synthetic negatives. On the TPpred-LE benchmark it achieves 55.3% Micro and 38.6% Macro F1, on par with the prior state of the art (57.9% Micro F1 / 38.1% Macro F1). These numbers were obtained after removing all sequences overlapping the TPpred-LE test set to ensure a fair comparison.

Finally, an L1-regularized variant demonstrates that this performance need not come at the expense of interpretability: individual filters specialize into recognizable biological motif detectors—for example, the GTFT glucagon-like signature for lipid-metabolism peptides— learned without explicit motif supervision. Together, these results show that compact, interpretable convolutional models are a practical and trustworthy foundation for therapeutic-peptide screening.

## Acknowledgments

The authors thank David Pekker for fruitful conversations, Katsiaryna Tsarova for project coordination, and Cedric Kring for preparing the release version of the software.

## 7 Code, Model, and Data Availability

Code, trained model parameters, and the processed train, validation, and test splits used to reproduce the results will be made publicly available at https://github.com/terra-quantum-public/tq-therapep-ai upon public posting of this preprint. The repository includes instructions for preprocessing the data, regenerating synthetic decoys, and retraining the models.

## 8 Author Contributions

S.M. conceived the original idea for the study. R.E. developed the modeling approach, performed the experiments, analyzed the results, and wrote the initial manuscript. A.V., A.C.P., S.M., and M.R.P. contributed to study design, interpretation of results, and manuscript revision. All authors reviewed and approved the final manuscript.

## A Appendix

In the following, we show results comparing different architectural choices that substantiate the design choice of our final model in the main work. In Sec. A.1, we show a comparison of different architectures with comparable hyper-parameters. In Sec. A.2, we show a systematic scan of hyper-parameters. In Sec. A.3 we show Learning curves related to our early stopping choice. Lastly, in Sec. A.4, we show the label mapping for the external benchmark evaluation on the TPpred-LE benchmark.

### A.1 Architecture Comparison

We compared several neural network architectures under identical training conditions to understand which design choices are most effective for therapeutic peptide classification. Table 3 summarizes the results.

**Table 3:** Architecture Comparison. Test set performance on 49-label dataset. The models were trained with identical hyperparameters where applicable. A Random Forest (RF) model is shown for a baseline comparison.

| Model | Micro F1 | Macro F1 |
| --- | --- | --- |
| RF | 42.9 % | 20.9 % |
| CNN (parallel kernels) | 65.4% | 42.3% |
| CNN-Sequential | 67.7% | 42.5% |
| CNN-Dilated | 64.3% | 42.0% |
| CNN-Attention | 59.1% | 36.1% |
| ResNet | 66.1% | 41.6% |
| Transformer (TPpred-LE style) | 50.6% | 21.8% |
| **CNN (5× ensemble, fine-tuned)** | **78.9%** | **54.4%** |

The baseline model is a Random Forest (RF) with a performance of 42.9% Micro F1 and 20.9% Macro F1 which is significantly lower than all other models. We tested different CNN architectures with a. parallel kernels, b. sequential CNN kernels, and c. dilated kernels which show a significantly higher performance of 64.3%–67.7% Micro F1 and 42.0%–42.5% Macro F1 which we attribute to including translational symmetry. The variations within the classes are relatively low and we chose a combination of a 2-layer CNN with multiple kernels for the main work, leaving out dilated kernels.

A CNN model extended by an attention head performed worse than the original CNN model. Residual nets show strong performance in deep neural network architectures and can avoid vanishing gradient problems. We implemented a ResNet architecture which showed a performance close to the CNN architectures but overall, the CNN architectures slightly outperformed the ResNet architecture in both Micro- and Macro F1.

We implemented a Transformer encoder-decoder architecture closely following TPpred-LE, with label embeddings, function-function self-attention, function-residue cross-attention, and function-specific classifiers. Using the same training data and encoding (BLOSUM62), this architecture achieves 50.6% Micro F1 and 21.8% Macro F1, while the CNN variants achieve 65–68% Micro F1 on the same data.

One possible explanation for the strong performance of the CNN models is that therapeutic peptide classification relies primarily on local sequence patterns—such as amphipathic motifs in antimicrobial peptides or cysteine patterns in disulfide-rich toxins—which convolutional filters are well-suited to capture. Particularly when adding augmented negative data, motif detection becomes an important property which CNNs excel at due to their built-in locality. Our final model uses a 5-model CNN ensemble with parallel multi-scale kernels (sizes 3, 5, 7, 9), achieving 79.0% Micro F1 with approximately 15M total parameters.

### A.2 Hyperparameter Study

Table 4 summarizes the hyperparameter sensitivity analysis across all dimensions tested.

**Table 4:** Hyperparameter Study Results. Test set performance for different configurations. All experiments use CNN with enhanced encoding unless otherwise noted.

| Configuration | Parameters | Micro F1 (%) | Macro F1 (%) |
| --- | --- | --- | --- |
| *FC Dimensions* |  |  |  |
| 128→64 | — | 65.5 | 39.7 |
| 256→128 | — | 65.5 | 42.8 |
| 512→256 | — | 66.6 | 44.1 |
| 768→384 | — | 67.5 | 45.4 |
| *Dropout* |  |  |  |
| p=0.1 | — | 70.0 | 46.7 |
| p=0.2 | — | 68.1 | 43.5 |
| p=0.3 | — | 65.2 | 42.4 |
| p=0.4 | — | 64.4 | 38.9 |
| p=0.5 | — | 62.0 | 34.9 |
| *Learning Rate* |  |  |  |
| lr=0.0001 | — | 65.1 | 39.1 |
| lr=0.0005 | — | 64.8 | 38.3 |
| lr=0.001 | — | 66.2 | 42.5 |
| lr=0.002 | — | 64.9 | 42.4 |
| *Ensemble Size* |  |  |  |
| 1 (single) | ~1M | 65.9 | 42.2 |
| 3 models | ~3M | 67.6 | 44.0 |
| 5 models | ~5M | 68.9 | **46.2** |
| 10 models | ~10M | **69.0** | 45.5 |
| *Pooling Strategy* |  |  |  |
| Max (baseline) | ~1.0M | 65.4 | 42.3 |
| Max + Mean | ~1.1M | 65.6 | 42.2 |
| *Encoding* |  |  |  |
| One-Hot | — | 65.9 | 43.1 |
| Enhanced (One-Hot + 14d Physical) | — | 65.5 | 41.3 |
| Learned (64d) | — | 65.5 | 42.4 |
| Learned (128d) | — | 65.1 | 41.6 |

Key findings:

- **FC Dimensions**: Larger hidden layers (768 →384) improve both Micro F1 (+2%) and Macro F1 (+5.7%) over smaller configurations.
- **Dropout**: Lower dropout (p=0.1) yields the best performance (70.0% Micro F1, 46.7% Macro F1), with performance degrading as dropout increases.
- **Learning Rate**: The default lr=0.001 performs well; both higher and lower values reduce performance.
- **Ensemble Size**: The 5-model ensemble achieves the best Macro F1 (46.2%) and near-optimal Micro F1 (68.9%), with diminishing returns beyond 5 models. Even with 5 models, our ensemble (5M parameters) remains substantially smaller than TPpred-LE (23M parameters). Note that the final finetuned model is larger (15M).
- **Pooling Strategy**: No meaningful difference between max pooling and max+mean pooling, suggesting global max pooling already captures essential discriminative features.

After systematic tests, we increased the model size and iteratively added slight improvements to the architecture to arrive at a final finetuned model.

### A.3 Learning Curves

Figure 10 shows training dynamics for the baseline CNN model. The model converges within 50 epochs, with validation loss plateauing around epoch 25. The gap between training and validation F1 indicates some overfitting, controlled by early stopping. Micro F1 and Macro F1 consistently kept increasing long after validation loss started increasing again. Therefore, we used the validation set’s Micro and Macro F1 as the early stopping criterion with a patience of 10 epochs as depicted in the Figure.

**Figure 10:**
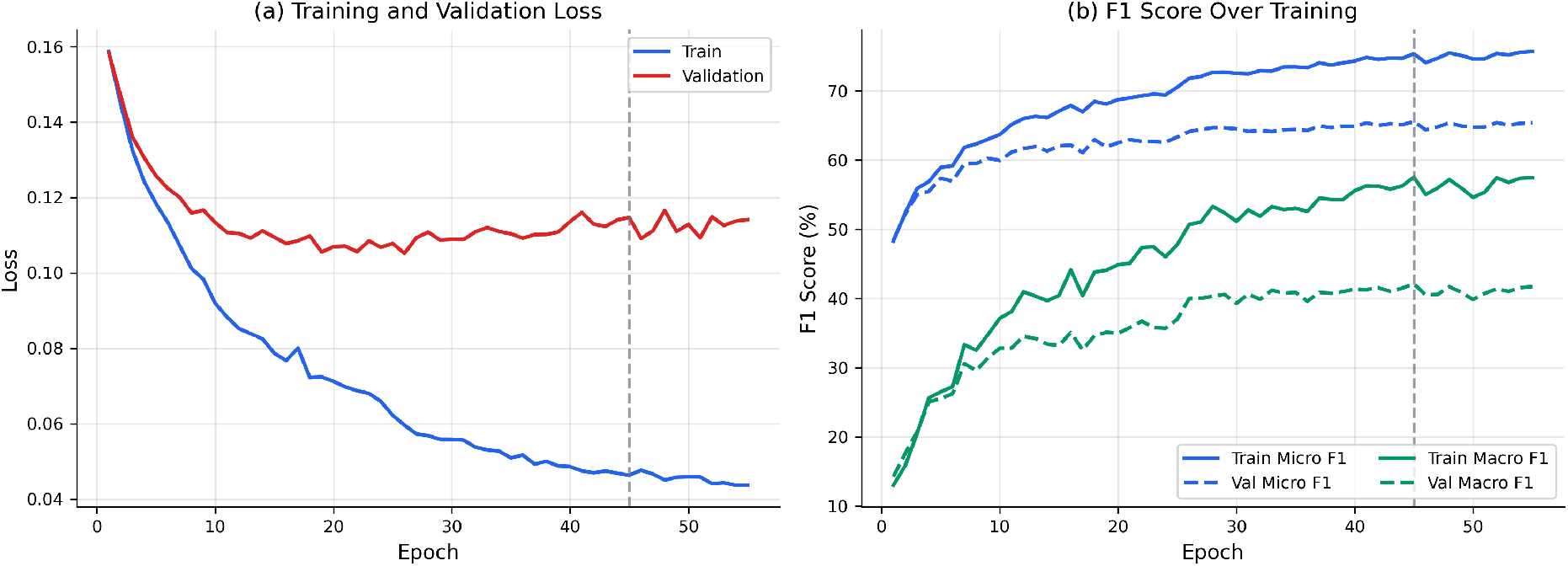
Training Dynamics. (a) Training and validation loss over epochs. (b) Micro F1 and Macro F1 for training and validation sets. The vertical dashed line indicates the best epoch.

### A.4 Label mapping between TheraPep-AI (49 labels) and TPpred-LE (15 labels)

TPpred-LE’s benchmark defines 15 therapeutic-peptide categories that are typically broader than the labels used in TheraPepDB. For example, TPpred-LE’s *AMP* (antimicrobial peptide) is an umbrella category that aggregates several finer subclasses in our taxonomy (*Antibacterial, Antifungal, Antiparasitic*, gram-specific antibacterial peptides, and the four antiprotozoal sub-labels). To compare the two models on a common label space we defined a semantic mapping from each TPpred-LE label to the set of TheraPep-AI labels it subsumes (Table 5). At evaluation time, the prediction for a TPpred-LE label is taken to be positive if any of its mapped TheraPep-AI labels is predicted positive (logical OR); the same rule is applied to the groundtruth labels. Of the 15 TPpred-LE categories, 12 admit an unambiguous semantic counterpart. Three labels are excluded from the comparison: *TXP* (toxic peptides) and *CCC* (cell–cell communication) have no direct equivalent in TheraPepDB, and *PBP* (penetrating brain peptides) is not separately annotated in our source. Metrics in Section 3.2 are then computed only over the resulting 12-dimensional subspace. This is a conservative setting for our model, which is trained to predict the full 49-label space yet is only credited for the 12 shared categories.

**Table 5:** Mapping between TPpred-LE labels (15 total) and TheraPep-AI labels (49 total). A TPpred-LE label is considered positive iff at least one of the mapped TheraPep-AI labels is positive (logical OR). The bottom three rows list TPpred-LE categories that are excluded from the cross-model comparison because they have no semantic counterpart in TheraPepDB.

| TPpred-LE | Description | Corresponding TheraPep-AI labels |
| --- | --- | --- |
| AMP | Antimicrobial | Antimicrobial, Antibacterial, Antifungal, Antiparasitic, Anti-gram+, Anti-gram-, Antiplasmodial, Antileishmania, Antimalarial, Antitrypanosomic |
| ABP | Antibacterial | Antibacterial, Anti-gram+, Anti-gram- |
| AIP | Anti-inflammatory | Antiinflammatory |
| AVP | Antiviral | Antiviral, AntiHIV, AntiHCV, AntiHSV, AntiMERS-Cov, AntiSARS |
| ACP | Anticancer | Anticancer, AntiBreastcancer, AntiCervicalcancer, AntiColoncancer, AntiLivercancer, AntiLungcancer, AntiProstatecancer, AntiSkincancer |
| AFP | Antifungal | Antifungal |
| DDV | Drug delivery | Drug_delivery |
| CPP | Cell-penetrating | Cell_penetrating |
| APP | Antiparasitic | Antiparasitic, Antiplasmodial, Antileishmania, Antimalarial, Antitrypanosomic |
| AAP | Anti-angiogenic | AntiAngiogenesis |
| AHTP | Antihypertensive | Antihypertensive |
| QSP | Quorum-sensing | Quorum_sensing |
| *Excluded — no TheraPep-AI counterpart* |  |  |
| TXP | Toxic | — |
| CCC | Cell-cell comm. | — |
| PBP | Penetrating brain | — |

